# Winner(s)-take-all: nonlinear amplification of DNA-encoded library

**DOI:** 10.1101/744680

**Authors:** Meiying Cui, Francesco Reddavide, Stephan Heiden, Luca Mannocci, Michael Thompson, Yixin Zhang

## Abstract

Information processing functions are essential for biological organisms to perceive and react to their complex enviornment, as well as for human to analyze and rationalize them. While our brain has an extraordinary power to process complex information, winner(s)-take-all computation is one of the simplest models of lateral inhibition and competition among biological neurons. It has been implemented as DNA-based neural networks, for example, to mimic pattern recognition. However, the utility of DNA-based computation in information processing for real biotechnological applications remains to be demonstrated. In this paper, we developed a winner(s)-take-all method for non-linear amplification of mixtures of DNA sequences. Unlike conventional biological experiments, selected species were not directly subjected to analysis. Instead, parallel computation among myriad of different DNA sequences was carried out with a neural network-inspired winner-take-all function, to reduce the information complexity. The method could be used for various oligonucleotide-encoded libraries, as we have demonstrated its application in decoding and data analysis for selection experiment with DNA-encoded chemical library against protein target.

## Introduction

As the medium to encode genetic information, DNA has in recent years found many new applications, e.g. encoding chemical structures of combinatorial libraries, to form self-assembled nanostructures, computation and data storage^1–4^. However, different from the high-fidelity DNA synthesis by DNA polymerase in cell replication cycles, both chemical synthesis and sequencing of DNA are error-prone. For example, in the construction of DNA-encoded chemical library (DEL), mistakes can be caused by side reactions and low reactivity in the combinatorial chemical library synthesis (Figure **1a**, i), as well as in the encoding procedure (Fiure **1a**, ii)^5,6^. The affinity-based selection protocols with immobilized target proteins on polymer matrices may yield to decoding artifacts due to promiscuous interaction and protein misfolding (Figure **1a**, iii and iv). PCR and NGS steps can also cause remarkable biases, while the difference in GC content is the most known origin (Figure **1a**, v,vi,and vii)^7^. Indeed, the evaluation of the selection is typically accomplished by ranking the NGS-sequence counts and it is based on the assumption that the binding affinity of the detected SMC (small molecular compound) to the native target protein, the amount of SMC-DNA conjugate captured on the polymer matrix, the PCR amplification, and the NGS reads are linearly correlated (Figure **1a**). Unfortunately, errors and biases accumulate over the steps, and poor correlations between the measured binding affinities of selected compounds to target protein and the counts have frequently been observed^5,8^. DELs with more sophisticated structures have been constructed in recent years, exacerbating the problem by adding more synthetic and DNA-manipulation steps^9–13^. While the challenges are associated with individual chemical and biochemical reactions, people also started to use statistic tools in data analysis^14–16^. However, as another undesired consequence of increasing library size, NGS cannot provide sufficient sequencing depth for comprehensive statistical analysis.

**Figure 1.**
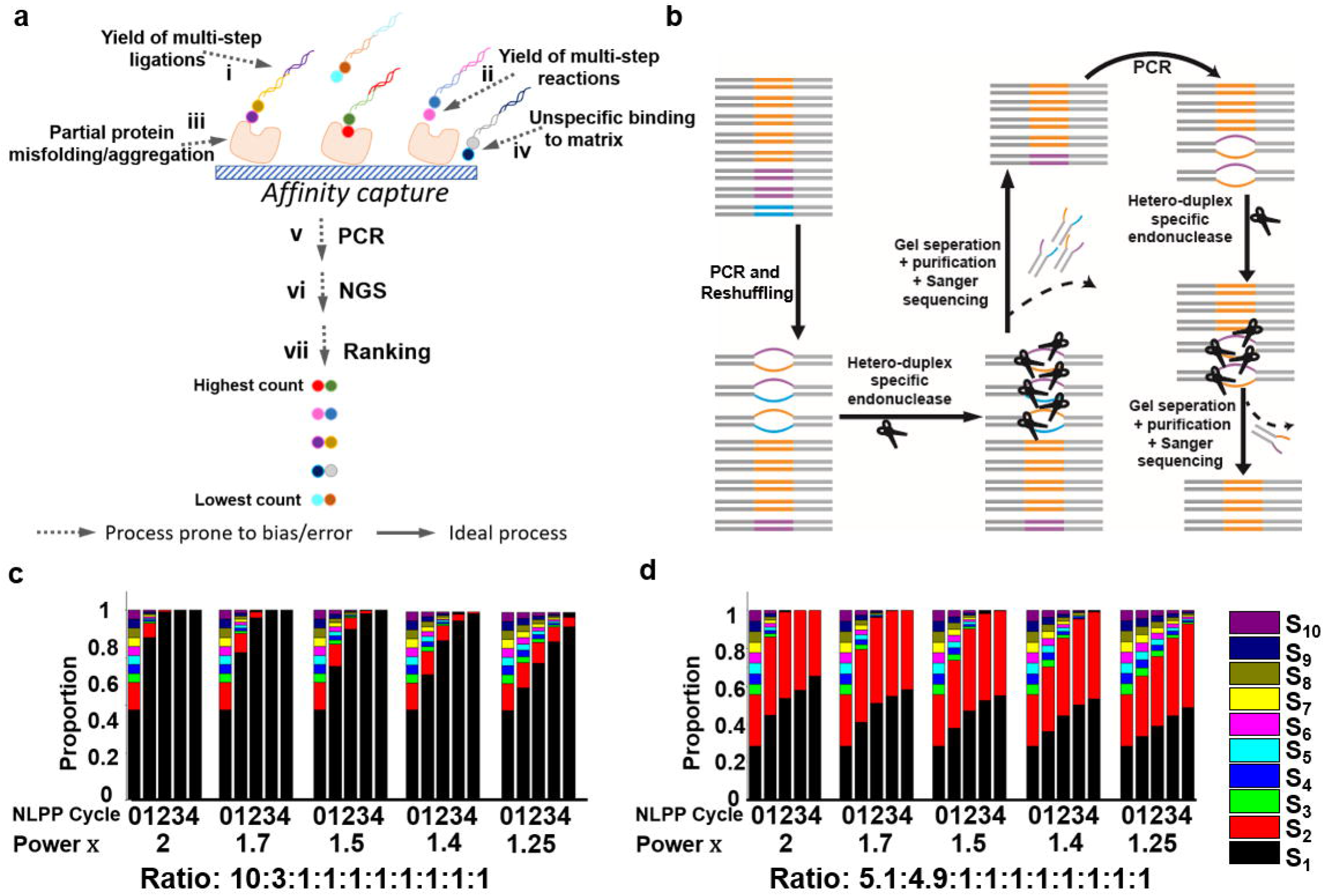
a) Compromised linearity in DEL experiment flow. If a selection experiment against target protein of interest is carried out at ideal condition using a perfect DEL, and NGS generates reads >> library size and there are no biases in PCR and NGS, the final ranking of sequence reads will be correlated with the binding affinities to protein for each chemical structure conjugated to its DNA barcode. Unfortunately, DEL experiments involve multiple steps, and each of them can undermine the potential linear correlation. b) Scheme of non-linear amplification of DEL (NADEL). The diverse DNA mixture is amplified via primers annealing to the constant region (grey lines). When primers are depleted, hetero-duplex DNAs are formed through code regions (orange, violet, and blue lines) and can be cleaved by hetero-duplex specific nuclease. After gel separation only full-length DNA will be amplified in the next cycle. Hetero-duplex DNA resulted from PCR is again cleaved by endonuclease and removed by gel separation. Iteration of the process leads to non-linear amplification of “winner” DNA (orange code). c,d) Simulated NADEL propagation of 10 sequence-DNA mixture with different NADEL powers (2,1.7,1.5,1.4, and 1.25).

While all these artifacts associated with DEL (Figure **1a**) compromise linearity, we ask whether the quest for linearity could be ultimately avoided. Systems with linearity is essential for many technologies developed by human, as it simplifies the ways how we collect, analyze, process, and respond to different signals. However, there are many other more complex processes which cannot be reduced to a linear model (e.g. drug screening using DEL). On the other hand, linearity is often not intrinsic in biology, while winner-take-all, and probably more often, winners-take-all are among the simplest and powerful competitive models for evolution, as well as for neural networks and recognition^17,18^. In this work, we aim to develop a bio-inspired winner(s)-take-all function for non-linear amplification of DNA-Encoded Library (NADEL). If the affinity captured SMC-DNA conjugates could be subjected to a non-linear procedure, through which only those most enriched member(s) can survive and be amplified, the downstream selection data analysis could be remarkably cheaper and faster, even possible with Sanger sequencing.

## Result

The conceptual workflow of the Non-linear Amplification of DEL (NADEL) is described in Figure **1b**. Reshuffling a mixture of double stranded DNA carrying a randomized sequence region at denaturation temperature results in the formation of heteroduplexes, which can be selectively cleaved by mismatch-specific endonucleases such as surveyor nuclease, a widely used enzyme for mutation detection and error correction in synthetic DNA with high specificity^19–22^. During the reshuffling process, the relatively more abundant sequences have higher probabilities to form homoduplexes, thus are prevented from cleavage. The homoduplexes, as the “winners” from this process, but not the fragments from the cleaved heteroduplexes can be further amplified by PCR. The homoduplexes enriched sample can be then either purified and sequenced, or subjected to next NADEL cycle to further enhance the content of winner(s). In order to build a molecular circuit to realize the non-linear amplification of DNA information and winner(s)-take-all function, we first simulated the amplification processes of various mixtures containing ten different DNA sequences (Figure **1** and Figure **S2**). The percentage of the member *j* (*10 ≥ j ≥ 1*) in the mixture after *n+1* cycles of treatments (*P_j,n+1_*) is expressed by the following formula:

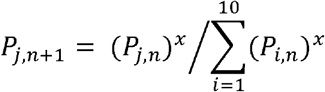

where *x* is a number between 1 and 2, reflecting the efficacy of the winner(s)-take-all process. For an NADEL cycle at ideal condition, *x* is 2, indicating all the mismatches generated by PCR are consumed by the nuclease, whereas for an ideal linear amplification (i.e., in absence of the nuclease), *x* is 1. As illustrated in the following experimental parts, performing NADEL cycles are relatively cumbersome. Therefore, we have limited the cycle number ≤ 4. For a mixture with only one member in large excess at the initial condition, even NADEL with low *x* value can completely eliminate the qPCR signal of the other members, though more cycles will be needed, as compared to NADEL with high *x* value (Figure **1c**). For a mixture with more than one major component (Figure **1d** and Figure **S2**), high *x* value is needed to eliminate the members of relatively low abundance. Moreover, depending on the initial ratios of different members, it could lead to more than one major species in the final mixture (winners-take-all).

To further test the experimental application of NADEL concepts, we designed a small library of 10 DNA sequences with codes of 20 nt (Figure **2a**), flanked by primers A and B. The library possesses high diversity as the difference between any two codes is > 15. The ten resulting sequences are mixed in 7 different ratios (Figure **2a**), mimicking various mixture compositions before and after DEL selections. For example, mixing ten sequences in equal amount (sample A_0_) represents an ideal library before selection. The sample B_0_ represents the situation of one strong binder with some weak binders, while the sample F_0_ mimics a selected mixture with multiple moderate binders. As a representative example, after one cycle of NADEL, sample A showed a remarkable decrease of the 60 bp band on the gel, as compared to the sample without treatment. The 60 bp DNA band was separated, purified and submitted to the next NADEL refining cycles.

**Figure 2.**
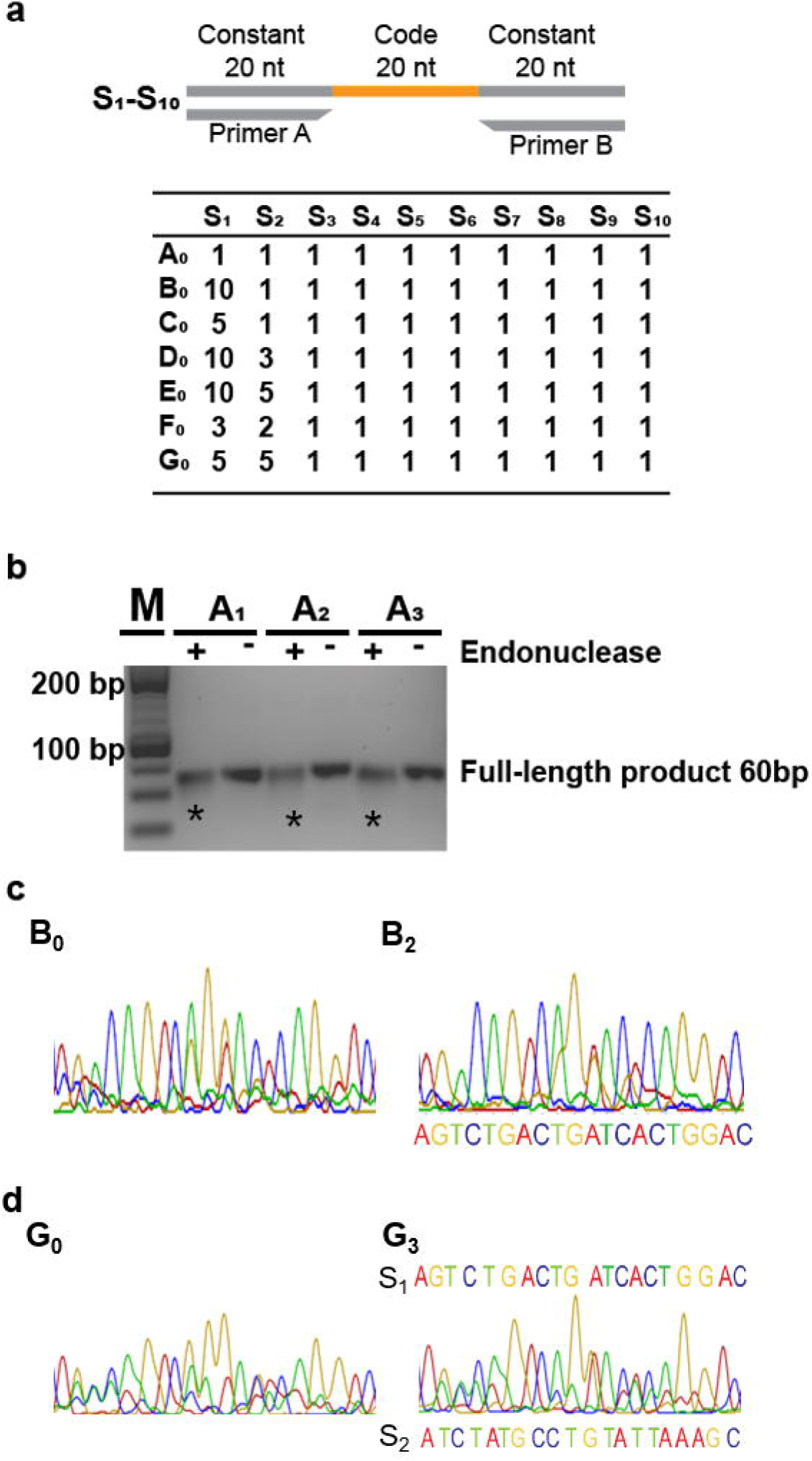
a) Scheme of 10 DNA sequences (S_1_-S_10_) constituting small libraries in 7 ratios (A_0_-G_0_). b) Representative agarose gel electrophoresis of sample A before (A_1−_), after one round (A_1+_, A_2−_), two rounds (A_2+_, A_3−_), and three rounds (A_3+_) of NADEL. Samples A_1−_, A_2−_, and A_3−_ are PCR products of A_0_, A_1+_, and A_2+_ after gel separation and purification, respectively. The bands of samples A_1+_, A_2+_, and A_3+_ show lower intensity compared to A_1−_, A_2−_, and A_3−_, respectively, and have smear below (*). M: marker c) Sanger sequencing chromatogram of sample before NADEL (B_0_) and after two rounds of NADEL (B_2_). d) Sanger Sequencing chromatogram of sample before NADEL (G_0_) and after three rounds of NADEL (G_3_).

In order to demonstrate the possibility of analyzing the NADEL products by Sanger sequencing, samples (B_0_ – G_0_) were subjected to NADEL cycles until the sequencing profile were dominated by the signals from the most abundant sequence with low ambiguity and high accuracy (Figure **2**). As expected, the NADEL sequencing profiles of A_0_ (A_1_, A_2_, and A_3_ where the subscript number is the number of NADEL cycles) are as noisy as the original untreated A_0_ sample (Figure **S3a**). For the other samples, in which the relative abundances of sequences are varied to mimic different selection results (sample B to G), the noise of sequencing profiles decreased after different NADEL cycles. For B_0_, C_0_, D_0_, and E_0_ samples, mimicking the presence of a dominantly enriched binder after the selection process, one to two NADEL cycles were sufficient to read the most abundant sequence **S_1_**without error (Figure **2c**, **2d**, and figure **S3**).

The more challenging samples are G_0_ and F_0_, in which two sequences (**S_1_** and **S_2_**) are in comparable excess (i.e., mimicking the presence of multiple equally strong binders after library selection). G_0_ could be particularly difficult, because both sequences are in equal amount. Interestingly, a single NADEL cycle of F_0_ led to the unambiguous identification of **S_1_**(Figure **S3b**). Differently, after the first NADEL cycle (G_1_, Figure S3f), the sequencing profile remained very noisy, while the noise has remarkably decreased following three NADEL cycles (G_3_, Figure **2d**). Eventually, reading the major peaks and the overlapping signals underneath, **S_1_** and **S_2_** could be both sequences fully retrieved (Figure **2d**). As conceptually anticipated, this result demonstrates that when there are more than one major species in relatively similar amount, NADEL cycles may lead to more than one major species in the final mixture (winners-take-all). As we will show later, when analyzing the samples from real selection experiments using NGS, multiple “winner” sequences can be co-evolved against other “loser” sequences. The experiments also allowed us to empirically estimate the *x* value above 1.4 for this 10-member library.

As DEL library are typically constructed comprising more coding domanins, we then tested the NADEL method with a DNA library containing two code regions (Figure **3a**). The code regions 1 and 2 contain 309 and 18 different sub-code sequences, 25 bp and 21 bp, respectively. The sub-codes are designed to ensure the difference between any pair > 8. The 5562-member library contains sequence **Q**, which accounts for 20% of the total DNA library mole (Figure **3a**). The non-linear amplification of **Q***vs.* the other 5561 sequences over NADEL cycles was simulated (Figure **3b**). Because of the additional constant domain between the two coding regions, the NADEL efficacy is expected to be lower than that for a single coding region.

**Figure 3.**
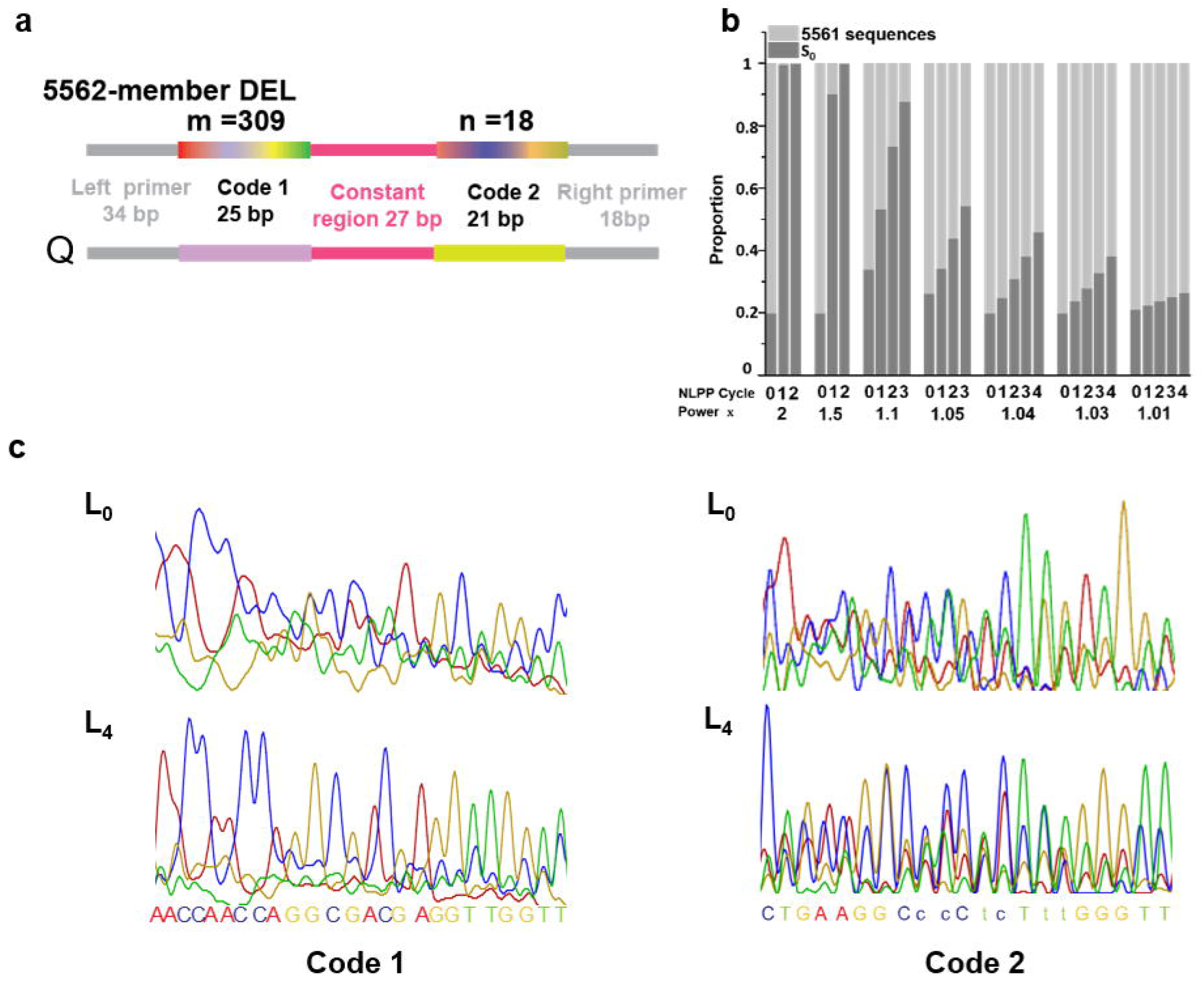
a) Scheme of 309×18-member library and sequence S_0_ with two code regions. b) Simulated sequence distribution of 5562 library with different NADEL powers after NADEL cycles. c) Sanger sequencing of code 1 and code 2 before NADEL and after four rounds of NADEL. Lowercase bases indicates the mismatches between Sanger sequencing read and actual code 2 of S_0_.

The mole ratio of sub-code 1 of **Q** to every other sub-code 1 sequence is 77:1; and the ratio of **Q** sub-code 2 to every other sub-code 2 sequence is 4.25:1. This design will allow us not only to test the NADEL method in a combinatorial library setup but also to evaluate its efficacy in simultaneously resolving coding regions on the same oligonucleotide with different randomization grade (e.g., sub-code 1 309-member *vs.* sub-code 2 18-member). Through the four NADEL cycles, the noise of sequencing profiles reduced gradually. Interestingly, after four cycles, the sequencing profile of code region 1 can be unambiguously assigned to the sub-code 1 of **Q** with zero error, while that of code region 2 contains six errors (Figure **3c**; errors in lower case). Neverthelss, the obtained sequence can still be assigned to the sub-code of **Q**, as they show the least hamming distance, as compared to the other library members (Table **S3**). The application of NADEL in decoding analysis is more powerful for sub-code region of large size, as the “winner(s)” can be more dominant over the other background noise sequences. It is important to note that by using different primers, NADEL can be performed either for individual code regions, or for the entire construct. NADEL efficacy for the entire construct is remarkably lower compared to single-code library, with *x* value over 1.05 according to the simulation. Nevertheless, whereas the use of NADEL for entire construct can reveal the most abundant combination after library selection, the application of NADEL for individual code regions can still rapidly and efficiently reveal the overall most enriched building blocks.

In order to test the utility of NADEL with a real DEL library, we performed selection experiment against protein target P using a DEL of 274 compounds with one code region of 20 nt (the identity of the target protein is not relevant to this study). The 274 DNA codes are designed to ensure the difference between any pair of sequences > 8. The NADEL cycles were followed by agarose gel electrophoresis (Figure **4a**). The sequencing profiles of the sample after selection P_0_, and the samples P_1_, P_2_, and P_3_ have shown a gradual reduction of noise over the NADEL cycles (Figure **4b**). Intriguingly, Sanger sequencing chromatogram could be deconvoluted into two coding sequences X and Y from major peaks and secondary signals respectively. To confirm this result, the samples were subjected to NGS (Illumina). In figure **4c** and **4d**, the NGS relative abundances for all codes were plotted for samples P_0_ – P_3_. While only a few enriched codes have shown a gradual increase of relative abundance, many codes that could be read in P_0_ cannot be detected in P_3_. Before NADEL treatment (P_0_), there is no code with zero read, whereas the numbers of codes with zero read in samples P_1_, P_2_, and P_3_ are 3, 6, and 15, respectively. As expected, the top two of the three codes which became dominant after NADEL cycles (Figure **4c**), were revealed to be X and Y, in good agreement with the Sanger sequencing result. Moreover, their relative abundances increased over the cycles, counting for more than 87 % of total reads after the third NADEL cycle, while the changes for code Z were small.

**Figure 4.**
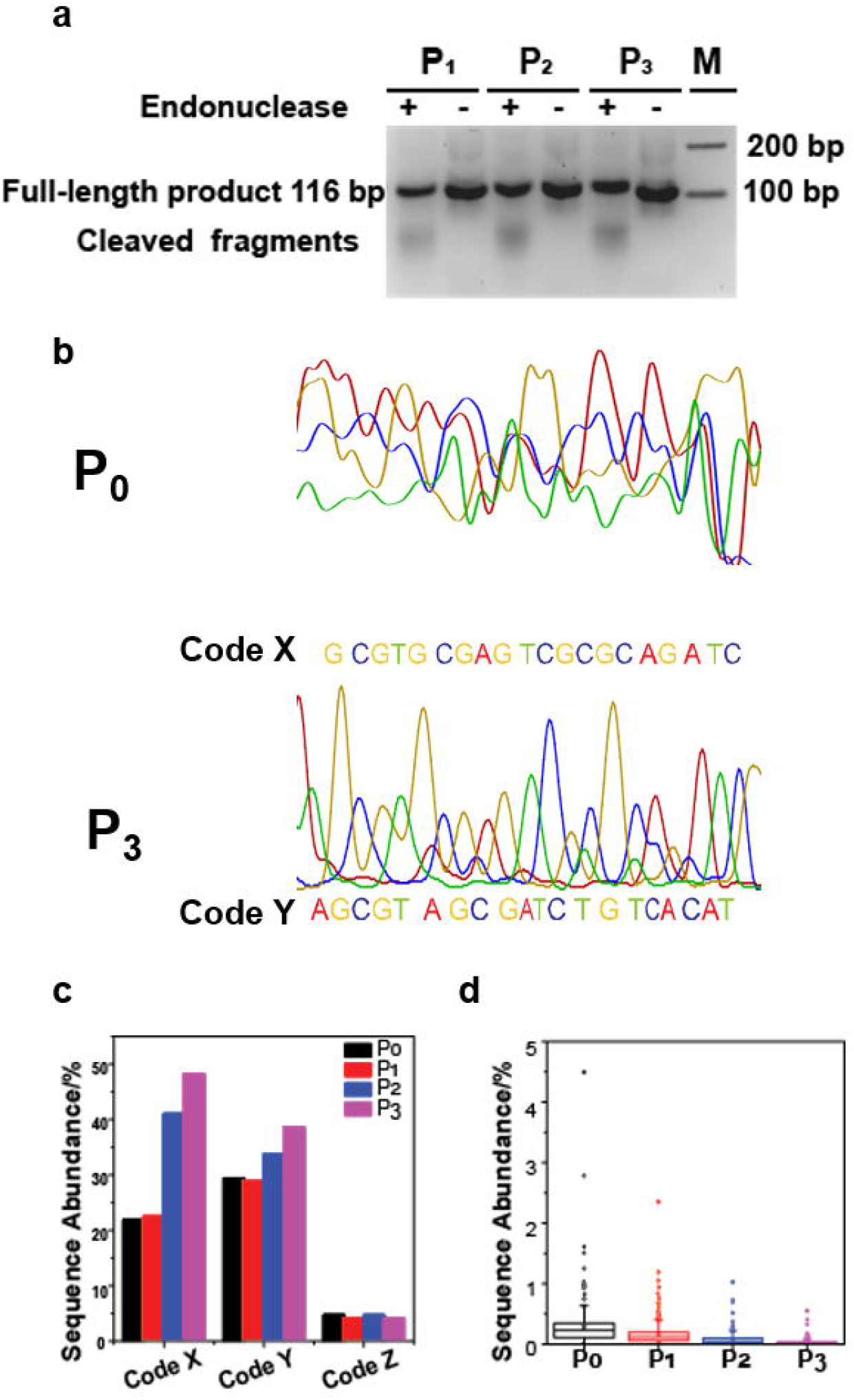
a) Agarose electrophoresis of selection output before (P_1−_), after one (P_1+_, P_2−_), two (P_2+_, P_3−_), three (P_3+_) rounds of NADEL. Samples P_1−_, P_2−_, P_3−_ are PCR products of P_0_, P_1+_, and P_2+_ after gel separation and purification, respectively. M: marker b) Sanger sequencing of selection output before NADEL and after three rounds of NADEL. c) Sequence abundance of code X, Y, and Z before and over three NADEL cycles. d) Boxplots representing sequence abundance of remaining 271 codes in NGS result from samples before and over three rounds of NADEL.

The NGS result also allows us to trace the fate of each code through the NADEL cycles. Except for codes X and Y, 269 codes have shown reduced relative abundance in P_3_ as compared to P_0_. Moreover, 204 out of the 269 codes have shown a step-by-step decrease through the entire process. This demonstrates that the non-linear process does not have a strong sequence bias.

## Discussion

With the NADEL approach, computation among different DNA sequences was carried out with a neural network-inspired winner-take-all function, to reduce the information complexity. Also different from traditional silicon-based computer technologies, the massive parallelism associated with DNA computing is especially attractive for huge DELs. DEL experiments involve multiple steps, and each of them can undermine the linear correlation between sequence counts and the number of affinity captured compounds by target protein (Figure **1a**). In selection experiment, enriched species are directly subjected to NGS analysis. However, with the tremendous increase of DEL size, it becomes increasingly difficult for NGS to provide sufficient sequencing depth allowing over-sampling and comprehensive statistical analysis. For example, the first NGS-decoded DEL has 4000 compounds in 2008, while DEL with 4 × 10^13^ compounds has been recently announced^1,23^. Although the non-linear information processing method cannot provide a direct solution to these problems, NADEL method allows reducing the complexity of the selection output and more confidently focusing on just a few potential candidates at the end of the selection campaign, which can be further validated by affinity measurements and inhibition assays. The selection experiment results shown in Figure **4** as well as the model samples E and G (Figure **2**) represent typical experimental situations observed in DEL screening, where multiple codes with counts above a cutoff are considered as candidates. By applying NADEL cycles, we demonstrate that it is possible to rapidly and conveniently narrow down the candidate list without losing key information.

The high cost of library synthesis and NGS, as well as the sophisticated decoding and data analysis have frequently hindered the wide application of DEL technology. Our results have demonstrated that it is possible to read the enriched sequences after NADEL cycles using Sanger sequencing. Therefore, while many laboratories can perform peptide/protein display or SELEX experiments by means of cost-effective and fast Sanger sequencing, NADEL offers a simple alternative for DEL decoding and data analysis. At the cost of using relatively long sequences to encode chemical structures, deconvolution of sequencing data can reveal multiple codes, especially when a retrievable algorithm is applied in code design^24^.

DEL technology will advance through improving library synthesis, or by suppressing the promiscuous interactions using in-solution or on-cell selection methodology^25–27^, or by implementing new techniques in decoding and data analysis. The winner(s)-take-all approach does not aim to replace these efforts to improve the linearity of DEL. However, by mimicking the lateral inhibition and competition among biological neurons in information processing, the winner(s)-take-all function can provide us with an alternative perspective for future DEL data analysis. Moreover, the non-linear amplification method can also be used to build molecular circuits to accomplish other sophisticated tasks. Recently Cherry et al. developed a two-species winner-take-all reaction, which has been successfully used to classify 10 × 10 patterns of handwritten digits ‘1’ to ‘9’^28^. NADEL can extend the two-species system to multiplex systems of eventually any number, allowing for the construction of neural networks reflecting the complexity of real world, while many NADEL experiments can be performed in parallel. Since manual NADEL implementation remain relatively cumbersome, as 4 cycles need 2 days, future automation will be very useful both to accelerate NADEL-based decoding and data analysis, as well as for building more sophisticated neural networks.

## References

1. Mannocci, L. et al. High-throughput sequencing allows the identification of binding molecules isolated from DNA-encoded chemical libraries. Proc. Natl. Acad. Sci. U. S. A. 105, 17670–17675 (2008).

2. Seelig, G. et al Enzyme-Free Nucleic Acid. 314, 1585–1589 (2006).

3. Rothemund, P. W. K. Folding DNA to create nanoscale shapes and patterns. Nature 440, 297–302 (2006).

4. Organick, L. et al. Random access in large-scale DNA data storage. Nat. Biotechnol. 36, 242–248 (2018).

5. Satz, A. L., Hochstrasser, R. & Petersen, A. C. Analysis of Current DNA Encoded Library Screening Data Indicates Higher False Negative Rates for Numerically Larger Libraries. ACS Comb. Sci. 19, 234–238 (2017)

6. Satz, A. L. Simulated Screens of DNA Encoded Libraries; the Potential Influence of Chemical Synthesis Fidelity on Interpretation of Structure-Activity-Relationships. ACS Comb. Sci. 18, 415–424 (2016).

7. Li, G. et al. Design, preparation, and selection of DNA-encoded dynamic libraries. Chem. Sci. 6, 7097–7104 (2015).

8. Yang, H. et al. Discovery of a potent class of PI3Kα inhibitors with unique binding mode via encoded library technology (ELT4). ACS Med. Chem. Lett. 6, 531–536 (2015).

9. Li, Y. et al. Versatile protein recognition by the encoded display of multiple chemical elements on a constant macrocyclic scaffold. Nat. Chem. 10, 441–448 (2018).

10. Stress, C., Sauter, B., Schneider, L., Sharpe, T. & Gillingham, D. A DNA-encoded chemical library incorporating elements of natural macrocycles. Angew. Chemie Int. Ed. 58, 9570–9574 (2019).

11. Lerner, R. A. et al. Functionality independent DNA encoding of complex natural products. Angew. Chemie Int. Ed. 131, 9355–9362 (2019)

12. Gerry, C. J., Wawer, M. J., Clemons, P. A. & Schreiber, S. L. DNA barcoding a complete matrix of stereoisomeric small molecules. J. Am. Chem. Soc. 141, 10225–10235 (2019)

13. Zhu, Z. et al. Design and Application of a DNA-Encoded Macrocyclic Peptide Library. ACS Chem. Biol. 13, 53–59 (2018)

14. Buller, F. et al. Discovery of TNF Inhibitors from a DNA-Encoded Chemical Library based on Diels-Alder Cycloaddition. Chem. Biol. 16, 1075–1086 (2009).

15. Buller, F. et al. High-throughput sequencing for the identification of binding molecules from DNA-encoded chemical libraries. Bioorganic Med. Chem. Lett. 20, 4188–4192 (2010).

16. Satz, A. L. DNA Encoded Library Selections and Insights Provided by Computational Simulations. ACS Chem. Biol. 10, 2237–2245 (2015).

17. Mori, K., Nagao, H. & Yoshihara, Y. The olfactory bulb: Coding and processing of odor molecule information. Science 286, 711–715 (1999).

18. Kaski, S. & Kohonen, T. Winner-take-all networks for physiological models of competitive learning. Neural Networks 7, 973–984 (1994).

19. Aggeli, D. et al. Diff-seq: A high throughput sequencing-based mismatch detection assay for DNA variant enrichment and discovery. Nucleic Acids Res. 46, 42 (2018).

20. Saaem, I., Ma, S., Quan, J. & Tian, J. Error correction of microchip synthesized genes using Surveyor nuclease. Nucleic Acids Res. 40, 23 (2012).

21. Lubock, N. B., Zhang, D., Sidore, A. M., Church, G. M. & Kosuri, S. A systematic comparison of error correction enzymes by next-generation sequencing. Nucleic Acids Res. 45, 9206–9217 (2017).

22. Hughes, R. A., Miklos, A. E. & Ellington, A. D. Enrichment of error-free synthetic DNA sequences by CEL I nuclease. Curr. Protoc. Mol. Biol. 1, 1–10 (2012).

23. Halford, B. Breakthroughs with bar codes. C&EN Glob. Enterp. 95, 28–33 (2017).

24. Y. Zhang, Herrmann, Jana, Wieduwild, Robert, Boden, Annett, TECHNISCHE UNIVERSITAET DRESDEN (Dresden, DE), United States, 2016.

25. Zhao, P. et al. Selection of DNA-encoded small molecule libraries against unmodified and non-immobilized protein targets. Angew. Chemie Int. Ed. 53, 10056–10059 (2014).

26. Chan, A. I., McGregor, L. M., Jain, T., & Liu, D. R. Discovery of a Covalent Kinase Inhibitor from a DNA-Encoded Small-Molecule Library × Protein Library Selection. J. Am. Chem. Soc. 139, 10192–10195 (2017)

27. Wu, Z. et al. Cell-based selection expands the utility of DNA-encoded small-molecule library technology to cell surface drug targets: Identification of novel antagonists of the NK3 tachykinin receptor. ACS Comb. Sci. 17, 722–731 (2015)

28. Cherry, M. K. & Qian, L. Scaling up molecular pattern recognition with DNA-based winner-take-all neural networks. Nature 559, 370–376 (2018)

