## Supplementary information for "Winner(s)-take-all: nonlinear amplification of DNA-encoded library"

1.	Materials and methods

1.1	Reagents and oligonucleotides

All oligonucleotides were purchased from IBA life sciences (Göttingen, Germany) and Metabion (Steinkirchen, Germany) in High-Performance Liquid Chromatography (HPLC)-purified grade, molecular biology grade, Next Generation Sequencing (NGS) grade or on the Controlled Pore Glass (CPG) solid support according to applications. Surveyor Mutation Detection Kit was purchased from IDT DNA Technologies (Coralville, Iowa, USA). Building blocks of the libraries were purchased from Sigma-Aldrich (St. Louis, MO, USA), Enamine (Kiev, Ukraine), Alinda Chemical (Moscow, Russia), ChemBridge Corporation (San Diego, CA, USA) and Maybridge Chemical Company (Altrincham, UK). Other reagents were unless otherwise noted in the text, were purchased from Thermo Fisher Scientific (Waltham, MA, USA).

1.2	Polymerase chain reaction (PCR) in Non-linear PCR protocol (NLPP)

The PCR mixture (50 μL) for all samples contained 10 X High GC buffer, primers (500 nM), dNTP mix (0.2 mM), and Phusion high-fidelity polymerase (1U) and 30 ng of respective template DNA.

Samples A-G and 5562-member library were amplified using (Primer A and Primer B), and (Primer C and D), respectively, with the following cycling conditions in PCR thermocycler (VWR, USA): 45 s at 98 °C, 30 cycles of 30 s at 98 °C, 1 min at 55 °C, and 30 s at 72 °C, closing the cycle, final extension for 10 min at 72 °C, and storing at 4°C.

DNA-encoded chemical library selection output P_0_ was amplified using primers Primer E and Primer F with the cycling conditions: 30 s at 95 °C, then 5 cycles of: 30 s at 98 °C, 1 min at 57 °C, and 30 s at 72 °C, closing the cycle, 25 cycles of: 30 s at 98°C, 1 min at 64 °C, and 30 s at 72 °C, followed by 5 min at 72 °C, and store at 4°C. P_1_ and P_2_ were amplified with the following protocol: 45 s at 98 °C, 30 cycles of 30 s at 98 °C, 1 min at 64 °C, and 30 s at 72 °C, closing the cycle, final extension for 10 min at 72 °C, and storing at 4°C.

1.3	PCR for Sanger sequencing

Before subjecting to Sanger sequencing, samples A-G and 5562-member library were further amplified with primer pairs (Primer G and Primer H), and (Primer I and Primer J), respectively, which can extend the length of the amplicon and therefore, ensure the read quality of code region of Sanger sequencing. The PCR protocol was as following. The PCR mixture (50 μL) contained 10 X High GC buffer, primers (500 nM), dNTP mix (0.2 mM), and Phusion high-fidelity polymerase (1U) and 1 μL of respective template DNA. The cycling conditions are 30 s at 95 °C, then 2 cycles of: 30 s at 98 °C, 1 min at 55 °C, and 30 s at 72 °C, closing the cycle, 18 cycles of: 30 s at 98°C, 1 min at 65 °C, and 30 s at 72 °C, followed by 10 min at 72 °C, and storing at 4°C.

Sanger sequencing primers for sample A-G, 5562-member library, and selection output were Primer K, Primer L, and Primer M, respectively. Sanger sequencing was performed by Eurofins genomics, Germany.

1.4	Surveyor Nuclease Treatment

Before treatment PCR product was confirmed by gel electrophoresis by loading 2 μL on 2% agarose gel (Bio-Rad, USA). Then the remaining PCR product was mixed with 4.8 μL of MgCl_2_ (150 mM), 1 μL of Surveyor Enhancer S (SES), and 2 μL of Surveyor Nuclease S (SNS) and incubated at 42 °C for 1 h in the PCR thermocycler. The reaction was stopped by adding 1/10 volume of the stop solution.

1.5	Agarose gel electrophoresis, DNA purification, and quantification

10 μL of each reaction mixture was transferred to new tubes for gel imaging. 5 μL of each sample was mixed with 1 μL of 6x agarose loading dye and loaded on 2% gel. The agarose gel images were taken using ChemiDoc MP System with Image Lab (Bio-Rad, USA).

The rest were loaded on 2% agarose gel to separate full-length DNA from cleaved fragments. A constant voltage of 90 V was applied for 3 h. DNA bands were visualized by a UV transilluminator (UVP, Germany). The DNA bands of correct size in both treated and untreated samples were sliced out and subjected to gel purification by using Nucleospin Gel and PCR clean-up kit (Macherey-Nagel, Germany). The concentration of purified DNA was measured with Nanodrop 2000 Spectrophotometers. Purified DNA entered next rounds of Surveyor nuclease treatment as described above.

1.6	Illumina high-throughput sequencing

The untreated selection output, as well as the gel-purified, nuclease-treated samples, were again amplified to attach specific oligonucleotides required for high-throughput sequencing. PCR product was recovered by gel extraction. The primer pairs for samples P_0_, P_1_, P_2_, and P_3_ were (P_0_FOR, P_0_REV), (P_1_FOR, P_1_REV), (P_2_FOR, P_2_REV), and (P_3_FOR, P_3_REV), respectively. PCR protocol was the same as the protocol for Sanger sequencing sample preparation.

The concentration of each PCR product was measured by Nanodrop 2000 spectrophotometers and four samples were mixed in the same concentration and subjected to the 2nd PCR amplification with the primers Illumina FOR and Illumina REV to attach adapter sequences compatible with the flow cell. Final amplification product, which contained information of all four samples was subjected to high-throughput sequencing. High-throughput sequencing was performed with HiSeq2000 Next Generation Sequencer (Illumina, USA). Data analysis was performed following a previous report.^[1]^

1.7	Synthesis of 274-member library with one code region

The 274-member library was prepared by conjugating each building block (as carboxylic acid) to C6 amino-modified DNA on a CPG solid support. 1 eq (0.01mmol, 50 mM) carboxylic acid, 1 eq (0.01 mmol, 50 mM) 1-[Bis(dimethylamino)methylene]-1H-1,2,3-triazolo[4,5-b]pyridinium 3-oxid hexafluorophosphate (HATU) and 1 eq (0.1mmol, 50 mM) 1-hydroxy-7-azabenzotriazole (HoAt) were dissolved in 200 μL dimethyl sulfoxide (DMSO) and stirred 30 min on orbital shaker at 300 rpm at room temperature. The reaction mixture was added on top of the CPG beads and 3 eq. (150 mM) of N,N-Diisopropylethylamine (DIPEA) was added and stirred overnight at room temperature, then washed with DMSO and methylene dichloride (DCM). The DNA was cleaved from the CPG beads and deprotected with 1:1 mixture of ammonium hydroxide and aqueous methyl amine (AMA), while stirring on an orbital shaker at 420 rpm for 3 h at room temperature. The AMA solution was evaporated from the crude conjugates at 50 °C. The crudes were re-suspended with 1 mL MilliQ water and purified via reverse-phase HPLC (Waters, USA) on a Clarity 3u Oligo-RT C18 reverse−phase HPLC column (Phenomenex, Torrance, CA, USA), applying a gradient from 5% acetonitrile (ACN) to 35% ACN over 30 min. The correct fraction was confirmed via ultra-performance liquid chromatography-electrospray tandem mass spectrometry (UPLC-ESI-MS, Waters, USA) equipped with an analytical ACQUITY UPLC OST C18 column (Waters, USA). The fractions were dried using a speedvac dryer (Christ RVC 2-25 CD plus/CT 02-50 SR, Osterode am Harz, Germany) equipped with a PC 3000 series vacuum controller (Vacuumbrand, Wertheim, Germany). According to the absorbance at 260 nm, the conjugates were quantified and equilibrated to 100 μM. Successfully coupled DNA-building blocks were then phosphorylated using T4 PNK (10 U, New England Biolabs, USA) in 1X NEB2 buffer supplemented with 1 mM ATP, in a total volume of 50 μL and encoded with respective code DNA sequences using T4 DNA ligase (4U, New England Biolabs, USA) in 1X NEB2 buffer supplemented with 1 mM ATP in 100 μL volume. The oligonucleotides were added in the following concentrations: 5 μM building block oligo-conjugate, 1.25 eq. of the respective code and ligation adapter. First, the oligonucleotides were heated at 60 ºC for 5 minutes and slowly cooled to 25 ºC over a time of 2 h to form dsDNA. Next, 4 U of T4 ligase and 1 μL of 100 mM ATP was added and incubated O/N at 18 ºC. Then, 1 µL (2 U) Klenow Fragment exo and 2 µL dNTP (2 mM) were added and incubated for 2 h at room temperature to form double-strand code. Both T4 DNA ligase and Klenow Fragment exo were inactivated by incubating at 75 ºC for 20 min and then slowly cooling to room temperature. Each ligation was tested via real-time quantitative PCR (qPCR, Thermo Fisher Scientific, USA) using the code flanking primers.

1.8	Synthesis of 309x18-member DNA library with two code regions

The 309 x 18-member DNA library was generated by ligating two sets of oligonucleotides. The first 309 oligonucleotides were 77 nt long containing 25 nt variable region, and the second set contained 18 oligonucleotides which were 48 nt long with 21 nt variable region. First set of oligonucleotides were pooled (each 1.5 nmol) and split into 18 tubes in equal amount. Then each phosphorylated oligonucleotide (each 556 nmol) from the second set was added in each tube of first set of oligonucleotides. Ligation was performed as descried in 1.7.

**1.9	Generation of 309x18 member library with 20% of S_0_**

Sequence S_0_ was generated by ligating two oligonucleotides, which were the two sets of oligonucleotides described in 1.8, respectively. Ligation of the oligonucleotides was performed in the same way as the encoding step in library synthesis.

Both ligation products (1.8 and 1.9) were PCR amplified and purified. The concentration of PCR products were then measured by Nanodrop 2000 Spectrophotometers. Then both samples were mixed accordingly to obtain a library, in which S_0_ account for 20% of the total mole.

1.10	Selection against target protein with 274-member library

The target protein (P) was immobilized on N-Hydroxysuccinimide (NHS)-activated Sepharose 4 Fast Flow (GE Healthcare, UK) via amide bond formation according to the manufacturer's instruction. The target protein on the beads was incubated with the library (1 nM per compound) for 1 h at room temperature. After washing the beads three times with 1x PBS buffer with 0.05% Tween-20, the bound library members were eluted in 100 μL 10 mM Tris buffer with 0.05% Tween-20, pH 8.3 by heat at 95 °C for 10 min.

2. Supplementary figures

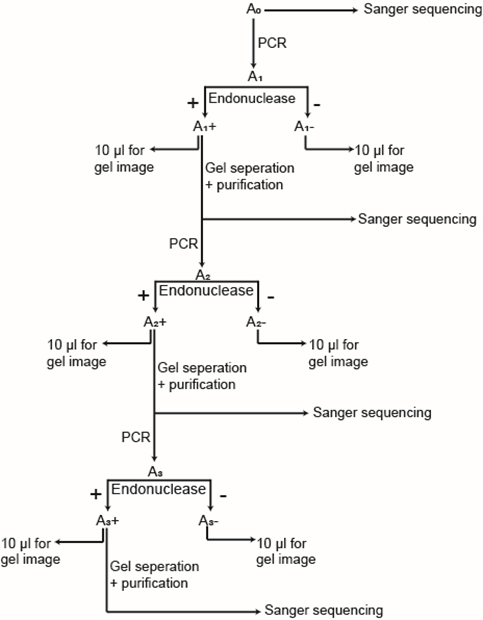

Figure S1. Detailed workflow of three-round NLPP (e.g. sample A0)

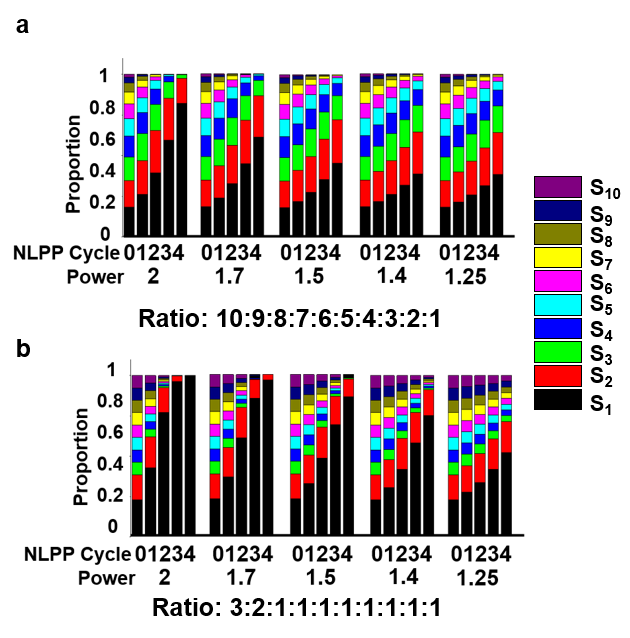

Figure S2.Simulated NLPP propagation of 10 sequence-DNA mixture with different NLPP powers (2,1.7,1.5,1.4, and 1.25).

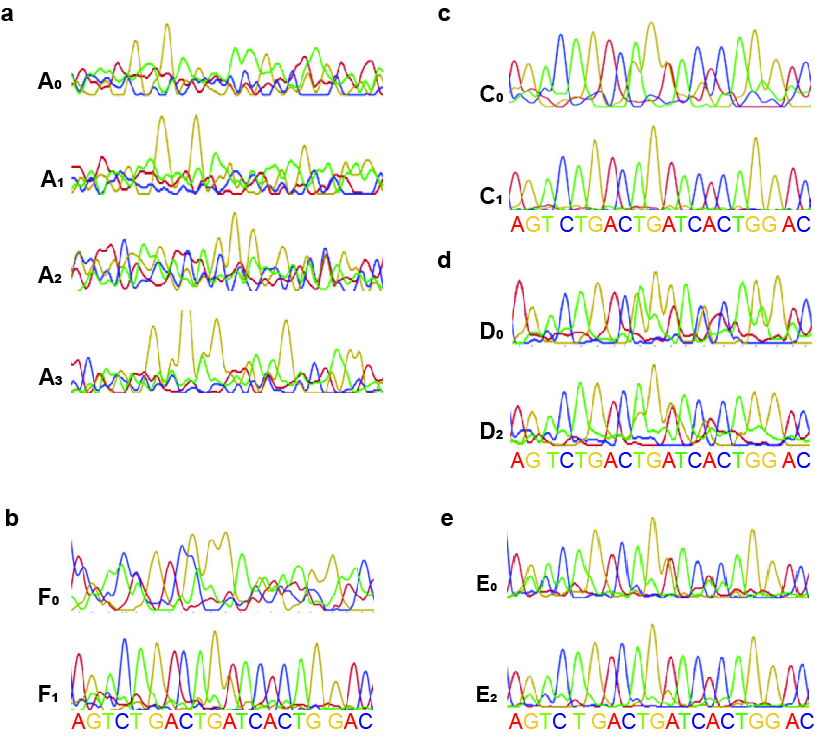

Figure S3. Sanger sequencing chromatogram of samples a) A_0_-A_3_, b) F_0_ and F_1_, c) C_0_ and C_1_, d) D_0_ andD_2_, and e) E_0_ and E_2_.

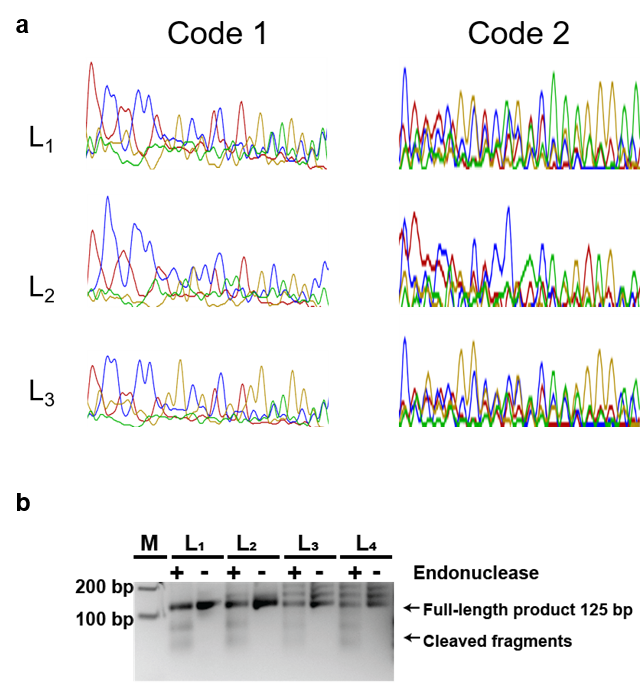

Figure S4. a)Sanger sequencing chromatogram of samples L1,L2, and L3. b)Agarose gel electrophoresis image of sample L0 before (L1-), after one round (L1+, L2-), two rounds (L2+, L3-), three rounds (L3+, L4-), and four rounds (L4+) of NLPP. Samples L1-, L2-, L3-, and L4- are PCR products of L0, L1+, L2+, and L3+ after gel separation and purification, respectively. M: 100 bp marker

3. Supplementary tables

**Table S1.** Sequence of 10 oligonucleotides comprising library samples A0-G0 (from 5’ to3’)

| Oligonucleotides | Sequence |
| --- | --- |
| S_1_ | GACAATTCACACACGTCCGCAGTCTGACTGATCACTGGACATGAGATCGGAAGAGCGTCG |
| S_2_ | GACAATTCACACACGTCCGCATCTATGCCTGTATTAAAGCATGAGATCGGAAGAGCGTCG |
| S_3_ | GACAATTCACACACGTCCGCACAAGGGGTCAAGCTCCTGTATGAGATCGGAAGAGCGTCG |
| S_4_ | GACAATTCACACACGTCCGCCCCTCCCTAAACTTTGCCTAATGAGATCGGAAGAGCGTCG |
| S_5_ | GACAATTCACACACGTCCGCGGGATCAGCCTGGGGATACAATGAGATCGGAAGAGCGTCG |
| S_6_ | GACAATTCACACACGTCCGCTATAAATACTACAAGCTCATATGAGATCGGAAGAGCGTCG |
| S_7_ | GACAATTCACACACGTCCGCGGGGCACCGCCAACAGAGATATGAGATCGGAAGAGCGTCG |
| S_8_ | GACAATTCACACACGTCCGCCCGGATTATATCCCATGAGGATGAGATCGGAAGAGCGTCG |
| S_9_ | GACAATTCACACACGTCCGCCTAGTGAAGTGATTCGTTCGATGAGATCGGAAGAGCGTCG |
| S_10_ | GACAATTCACACACGTCCGCGAACGTTAAGGGTGACCGTCATGAGATCGGAAGAGCGTCG |

Code regions are highlighted in blue

**Table S2.** Sequences of 309 x 18 oligonucleotides, S0 and Sequence X, Sequence Y, and Sequence Z (from 5’ to3’)

| Oligonucleotides | Sequence |
| --- | --- |
| First set (309) | GGAGGTGTAGACGACAGAGTATTTGACTGTCAGGNNNNNNNNNNNNNNNNNNNNNNNNNCCTTCTGGATTCGGTCGG |
| Second set (18) | AGCACCATCNNNNNNNNNNNNNNNNNNNNNACGCAATGTGGGAACAGC |
| S_0_ | GGAGGTGTAGACGACAGAGTATTTGACTGTCAGGAACCAACCAGGCGACGAGGTTGGTTCCTTCTGGATTCGGTCGGAGCACCATCCTGAAGGCGACGATCAGGGTTACGCAATGTGGGAACAGC |
| Sequence X | CAAGCAGAAGACGGCATACGAGATTGGTCTCAGCCGCCCTATGCGTGCGAGTCGCGCAGATCGTGGAGTTGCTCGATCTGGCTGAAGGCAGTAGGTTGTCGGGAGGGAAAGAGTGT |
| Sequence Y | CAAGCAGAAGACGGCATACGAGATTGGTCTCAGCCGCCCTATAGCGTAGCGACTGTCACATGTGGAGTTGCTCGATCTGGCTGAAGGCAGTAGGTTGTCGGGAGGGAAAGAGTGT |
| Sequence Z | CAAGCAGAAGACGGCATACGAGATTGGTCTCAGCCGCCCTATCTACACTATCGCTGCGTCACGTGGAGTTGCTCGATCTGGCTGAAGGCAGTAGGTTGTCGGGAGGGAAAGAGTGT |

Variable regions are highlighted in blue

**Table S3. Hamming distance between the measured sequence and library code 2 sequence**

|  | **Code 2** | **Hamming distance to the measured sequence** |
| --- | --- | --- |
| **S_0_** | **CTGAAGGCGACGATCAGGGTT** | **6** |
| **S_1_** | **CTTCTGCTGCAGAGAAGGGTT** | **12** |
| **S_2_** | **TTAAAGCTCGAGATTAAGGTT** | **10** |
| **S_3_** | **TTTGTATACGTACCAAAGGTT** | **14** |
| **S_4_** | **CAGTATTCGCAGTACTGGGTT** | **10** |
| **S_5_** | **AGCATGTCAGAGATGCTGGTT** | **13** |
| **S_6_** | **TAACTCGATCATGGTTAGGTT** | **12** |
| **S_7_** | **AAGACGCGATGACTCTTGGTT** | **11** |
| **S_8_** | **CAAACGGACTGACTTTGGGTT** | **7** |
| **S_9_** | **GAATTTGCGACTGATTCGGTT** | **11** |
| **S_10_** | **GGGCTTGACTGCAGCCCGGTT** | **14** |
| **S_11_** | **GGAAGGCTGATCATTCCGGTT** | **13** |
| **S_12_** | **AACCTGCGTCATCGGTTTAGT** | **15** |
| **S_13_** | **CGAGACCGTCATCCTCGAACC** | **14** |
| **S_14_** | **TCCAGCCTCTGACTGGAAACC** | **17** |
| **S_15_** | **TAACGTCTGCGCGGTTAAACC** | **18** |
| **S_16_** | **ACCGCAGCATCGACGGTAACC** | **18** |
| **S_17_** | **TTCTAATCGCATCAGAAAACC** | **15** |

**Table S4.** List of primers for PCR in NLPP and PCR for Sanger sequencing (from 5’ to3’)

| Primers | Sequence |
| --- | --- |
| Primer A | TATCAATCGTTACGGATACG |
| Primer B | CCTATGCAATCGACGCTCTT |
| Primer C | GCTGTTCCCACATTGCGT |
| Primer D | GGAGGTGTAGACGACAGAGT |
| Primer E | ACACTCTTTCCCTACCCGACAACCTACTGCCTTCAGCCAGATCGAGCAACTCC |
| Primer F | CAAGCAGAAGACGGCATACGAGATTGGTCTCAGCCGCCCTAT |
| Primer G | CAAGCAGAAGACGGCATACGAGATCGACGCTCTTCCGATCTCAT |
| Primer H | ACACTCTTTCCCTACCCGACAACCTACTGCCTTTCGAACCTAGACGAC AATTCACACACGTCCGC |
| Primer I | ACACTCTTTCCCTACCCGACAACCTACTGCCTTAAGCGGGCCCGTGCT  GTTCCCACATTGCGT |
| Primer J | CAAGCAGAAGACGGCATACGAGATGGAGGTGTAGACGACAGAGTATTTGAC |
| Primer K | ACACTCTTTCCCTACCCGACAACCTACTGCCTT |
| Primer L | CAAGCAGAAGACGGCATACGAGAT |
| Primer M | CAAGCAGAAGACGGCATACGAGAT |

**Table S5.** List of Illumina primers (from 5’ to3’)

| Primers | Sequence |
| --- | --- |
| P_0_FOR | ACACTCTTTCCCTACCCGACAACCTACTGCCTTCACACATCGAGCACACTCTTTCCCTCCCG |
| P_0_REV | CAAGCAGAAGACGGCATACGAGATTGGTCTCAGCCGCCCTAT |
| P_1_FOR | ACACTCTTTCCCTACCCGACAACCTACTGCCTTCTATGTAGCAGCACACTCTTTCCCTCCCG |
| P_1_REV | CAAGCAGAAGACGGCATACGAGATTGGTCTCAGCCGCCCTAT |
| P_2_FOR | ACACTCTTTCCCTACCCGACAACCTACTGCCTTCTCGTGATCATCACACTCTTTCCCTCCCG |
| P_2_REV | CAAGCAGAAGACGGCATACGAGATTGGTCTCAGCCGCCCTAT |
| P_3_FOR | ACACTCTTTCCCTACCCGACAACCTACTGCCTTCACACTAGACGAACACTCTTTCCCTCCCG |
| P_3_REV | CAAGCAGAAGACGGCATACGAGATTGGTCTCAGCCGCCCTAT |
| Illumina FOR | AATGATACGGCGACCACCGAGATCTACACTCTGAGCGATATACACTCTTTCCCTACCCGA |
| Illumina REV | CAAGCAGAAGACGGCATACGAGAT |

References

[1]	W. Decurtins, M. Wichert, R. M. Franzini, F. Buller, M. A. Stravs, Y. Zhang, D. Neri, J. Scheuermann, Nat. Protoc. 2016, 11, 764–780.

[2]	A. L. Satz, J. Cai, Y. Chen, R. Goodnow, F. Gruber, A. Kowalczyk, A. Petersen, G. Naderi-Oboodi, L. Orzechowski, Q. Strebel, Bioconjug. Chem. 2015, 26, 1623–1632.

[3]	D. R. Halpin, J. A. Lee, S. J. Wrenn, P. B. Harbury, PLoS Biol. 2004, 1031–1038.

Author Contributions

Y, Zhang and M. Cui designed the experiment. M. Cui conducted all NLPP experiment and selection experiment, F. Reddavide, and S. Heiden, M. Thompson synthesized DNA-encoded library. L. Mannocci, M. Cui, and Y. Zhang analyzed the data.
